# SLC16A6 is a tyrosine transporter for the melanosome

**DOI:** 10.64898/2026.07.06.736842

**Authors:** Corey N. Cunningham, Alex J. Bott, Charles H. Adelmann, Matthew Shields, Katarina E. Heyden, Jonathan G. Van Vranken, Álvaro Jesús Narbona-Pérez, Juan A. Cantres-Vélez, Guillaume Adelmant, Nathan M. Krah, Steven P. Gygi, David M. Sabatini, Jared Rutter

## Abstract

Cells enable specialized metabolism by compartmentalizing metabolic pathways into distinct organelles, which requires the membrane transport of metabolites. In melanocytes, the amino acid tyrosine is imported into developing melanosomes for the synthesis of the UV-protective pigment melanin^1,2^. In spite of extensive biochemical characterization, the identity of the melanosomal tyrosine transporter remains unknown. Here, we identify SLC16A6 as an orphan melanosome-localized metabolite transporter. Genetic screens reveal that *SLC16A6* expression is driven by the SOX10-MITF axis, the well-characterized master regulatory program governing melanogenesis and melanosomal homeostasis^3,4^. By redirecting SLC16A6 to the plasma membrane with an S240A mutation^5^, we demonstrate that SLC16A6 transports tyrosine, a process competitively inhibited by other bulky amino acids. We further determine that SLC16A6 is sufficient for *in vitro* melanosomal tyrosine uptake. Genetic depletion of *SLC16A6* triggered loss of melanosome biogenesis and function as well as depletion of most melanosomal components. Collectively, these findings establish SLC16A6 as a melanosomal tyrosine transporter that is essential for melanosome biogenesis.

## Main

Melanocytes synthesize the pigment melanin in specialized lysosome-related organelles called melanosomes^6^. These melanosomes are then ejected and taken up by keratinocytes, to be used for protection against UV irradiation. Within melanosomes, tyrosine is oxidized to L-DOPA by tyrosinase and subsequently converted through a series of enzymatic and non-enzymatic reactions to form eumelanins (derived exclusively from tyrosine) or pheomelanins (tyrosine- and cysteine-derived) (Fig. 1a)^7^. For complete melanin biogenesis, both tyrosine and cysteine must be imported into immature melanosomes. MFSD12 was recently identified as the melanosomal cysteine/cystine transport system^8^; however, the tyrosine transporter has remained elusive (Fig. 1a)^2^. The identity of the melanosomal tyrosine transport system has long been debated, but definitive biochemical evidence has ruled out previous candidates. For example, OCA2/P-protein was implicated in tyrosine transport because defects in this protein leads to oculocutaneous albinism^9,10^. However, recent functional studies have shown OCA2 transports chloride, which modulates pigmentation via melanosomal pH regulation^11,12^. With other candidates ruled out, the identity of the melanosomal tyrosine transporter has remained an open question. We reasoned that a tyrosine transporter for the melanosome would match the following criteria: (1) localization to early melanosomes; (2) classification within a solute carrier (SLC) subfamily with documented capacity for aromatic amino acid transport; and (3) strong co-expression with the melanosomal transcriptome, as evidenced by enrichment in pigmentation gene ontology terms among the most correlated genes.

**Figure 1.**
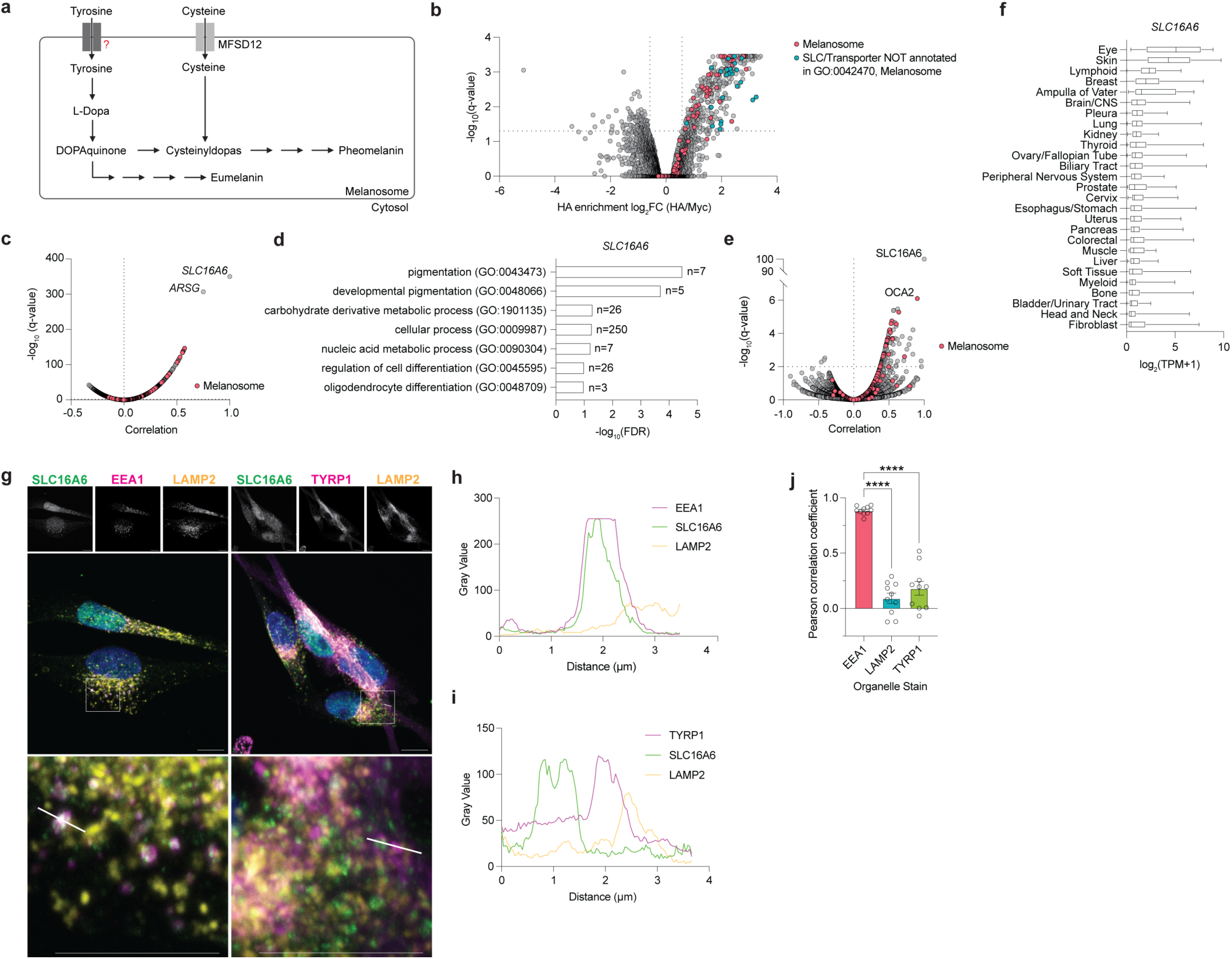
SLC16A6 is an orphaned early melanosomal transporter. **(a)** Illustration of melanogenesis. Tyrosine and cysteine are imported into developing melanosomes and converted to eumelanins and pheomelanins. **(b)** Volcano plot of proteins enriched in 3x-HA MelanoIP compared to control 3x-Myc MelanoIP from MeWo melanoma cells. **(c)** *SLC16A6* transcriptome-wide correlation. Co-expression analysis of *SLC16A6* transcripts versus all other transcripts using the DepMap short-read expression dataset (26Q1). **(d)** Panther GO-slim biological process analysis of top 500 most positively correlating transcripts from (c). **(e)** SLC16A6 proteome-wide correlation. Co-variation analysis of SLC16A6 versus all other proteins using the Harmonized MS CCLE Gygi dataset **(f)** *SLC16A6* expression (short-read) 26Q1 across various lineages. Data extracted from the Cancer Dependency Map. **(g)** Representative confocal microscopy images of endogenous SLC16A6 colocalization (50 cells per colocalization; scale bar 10 µm, n=3 biological replicates). **(h)** Representative line scan analysis of endogenous SLC16A6 colocalization with EEA1 and LAMP2; line is indicated in (g), left image. **(i)** Same as (h) but colocalization with TYRP1 and LAMP2; line is indicated in (g), right image. **(j)** Pearson correlation coefficient analysis of line scan averages from (g). 10 cells analyzed per condition with 5 random line scan analyses performed per cell. Average Pearson correlation coefficients per cell plotted. (\*\*\*\**P* < 0.0001).

### SLC16A6 localizes to melanosomes

To generate a candidate set of plausible tyrosine transporters localized to the melanosome, we used MelanoIP, a rapid melanosomal purification method^8^, followed by proteomics (Fig. 1b). This yielded a selective list of putative transporters, which we further prioritized based on localization^13,14^ and membership in SLC families with known aromatic amino acid substrates^15^ (Supplementary Table 1). Among these, SLC16A6 and SLC49A3 emerged as potential candidates. However, we deprioritized *SLC49A3* owing to its poor association with melanosomal genes (Fig. S1a,b). In contrast, *SLC16A6* expression positively correlated with the melanosomal gene network and pigmentation (Fig. 1c,d). These data were further corroborated by SLC16A6 protein co-variation, which showed the strongest positive correlation with the melanosomal chloride transporter OCA2/P-protein (Fig. 1e).

SLC16A6 is a poorly defined 12-pass transmembrane solute carrier with high expression in cutaneous and uveal melanoma cell lines, mature melanocytes, and the retina, which is also pigmented (Fig. 1f, Fig. S1c,d, Supplementary Table 1)^16,17^. SLC16A6 belongs to the SLC16A subfamily, whose members are proton-linked monocarboxylate transporters^18^. Of note, two members, SLC16A10 and SLC16A2, can transport aromatic amino acids and their derivatives (Supplementary Table 1)^19,20^. Given these findings, we sought to examine whether SLC16A6 indeed localizes to the early melanosomal membrane system.

We performed confocal microscopy in various melanoma cell lines of both endogenous and ectopically expressed SLC16A6. First, we observed that endogenous SLC16A6 co-localizes with early endosome antigen 1 (EEA1) (Fig. 1g-j). EEA1 is a marker of early endosomes from which later-stage melanosomes arise^6^. Previous studies have shown that the key melanosomal protein, PMEL, colocalizes with EEA1 in early-stage melanosomes^21,22^. Ectopic SLC16A6, which can be visualized with much more sensitivity due to use of the FLAG epitope/antibody, also localizes to late-stage melanosomes that contain enzymes and structural proteins such as TYRP1 and LAMP2^6,23,24^, suggesting that it can populate throughout the melanosomal pathway (Fig. S1e-h). Together, these results suggest that SLC16A6 is a melanosome-localized transporter.

### SOX10 and MITF drive SLC16A6 expression

Reasoning that its transcriptional regulators would likely be instructive as to its physiological function, we next sought to identify the transcriptional regulators of *SLC16A6* using Targeted Readout to Understand Transcription via Fluorescent *In Situ* Hybridization (TRoUT-FISH)^25^. Briefly, MeWo cells were transduced with a pooled library of CRISPR sgRNAs targeting human transcription factor and epigenetic genes, and we then performed RNA-FISH assessing endogenous *SLC16A6* mRNA abundance. We FACS-sorted cell populations with the highest and lowest *SLC16A6* abundance and sequenced their associated sgRNAs (Fig. 2a). We found that sgRNAs targeting *SOX10* and *MITF* were the two most enriched in cells with depleted *SLC16A6*, suggesting them as the primary transcriptional activators of *SLC16A6* (Fig. 2a,b). During mammalian development, SOX10 specifies and maintains the melanocyte lineage by binding to the *MITF* promoter driving its expression^3^. MITF activates its own expression as well as a transcriptional network responsible for melanosome biogenesis and maturation^3,4^.

**Figure 2.**
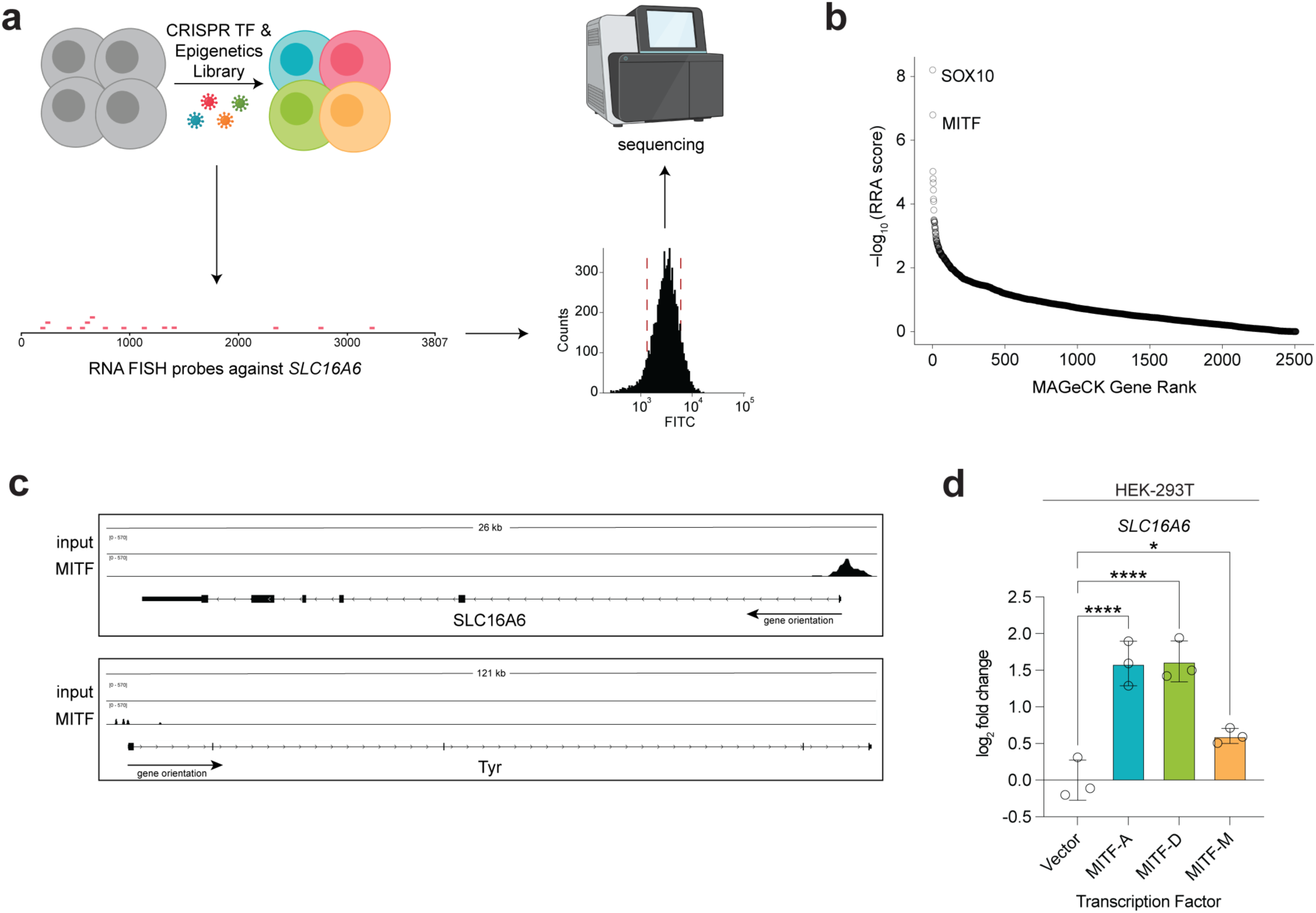
Master melanogenesis transcription factors, SOX10 and MITF, drive *SLC16A6* expression. **(a)** Schematic of TRoUT-FISH screen. Human MeWo melanoma cells were transduced with a pooled CRISPR library of the human epigenetic-focused sgRNA library, grown, puro-selected, and then fixed. RNA FISH targeting *SLC16A6* transcripts was performed followed by flow cytometry to isolate population of cells with lowest and highest fluorescence and then sequenced to determine sgRNA abundances. **(b)** TRoUT-FISH results of (a). RRA scores derived from MAGeCK comparing the high and low populations of *SLC16A6* abundance. **(c)** Genome browser tracks visualizing MITF chromatin-immunoprecipitation-sequencing (ChIP-seq) in MeWo KRAB cells^25^. **(d)** qPCR of *SLC16A6* after transient transfection of MITF isoforms in HEK-293T cells (\**P* < 0.05; \*\*\*\**P* < 0.0001).

ChIP-seq demonstrated that MITF directly binds to the *SLC16A6* promoter with an occupancy similar to that of known MITF target genes, such as tyrosinase (*TYR*), *SLC45A2*, and *DCT/TYRP2* (Fig 2c, Fig. S2a)^26^. In follow-up experiments, we overexpressed the three MITF isoforms in HEK-293T cells and each caused an upregulation of *SLC16A6* mRNA (Fig. 2d). Further supporting this regulatory link, a CRISPR-generated *MITF* mutant and RNAi-mediated *MITF* knockdown significantly downregulated *SLC16A6*, while MITF induction increased *SLC16A6* transcript levels (Fig. S2b-d)^27^. These results establish *SLC16A6* as a melanogenesis network gene driven by SOX10 and MITF.

### SLC16A6 mediates tyrosine transport

Although SLC16A6 belongs to the monocarboxylate transporter (SLC16A) family, its physiological substrates and functions have remained poorly defined. While prior reports suggested SLC16A6 acts as a plasma-membrane ketone body or low-affinity taurine transporter^28,29^, these roles are hard to rationalize with its intracellular localization and co-expression with melanosomal machinery. We therefore sought to assess the endogenous substrates of SLC16A6.

To facilitate an untargeted metabolomics-based screen for substrates, we sought to redirect the transporter to the cell surface using an SLC16A6 S240A mutant (Fig. 3a)^5^. The S240 residue of SLC16A6 is located within a large, 90-amino acid loop between transmembrane regions 6 and 7^30^, and was identified as critical for regulating protein turnover. Introducing this mutation (S240A) both extends the half-life of the transporter and, importantly, causes it to accumulate at the plasma membrane^5^. While both wildtype (WT) SLC16A6 and SLC16A6^S240A^ expression might change intracellular steady-state metabolite abundances, we predicted that SLC16A6^S240A^ would have the largest metabolomic change from the extracellular milieu compared to the intracellular space, given its plasma membrane localization. Therefore, we used expression of this mutant as a cell-based model of metabolite uptake, in a similar manner to how MFSD12-PM was used to study melanosomal cysteine uptake (Fig. 3b)^8^. Only in SLC16A6^S240A^-expressing HEK-293T cells, amino acids such as L-tyrosine and the branched-chain amino acids showed significantly decreased Δabundance (extracellular-intracellular), consistent with accelerated transport from the extracellular to intracellular metabolite pool (Fig. 3b, Fig. S3a). Consistent with this, L-tyrosine levels were significantly increased intracellularly and decreased extracellularly in SLC16A6^S240A^-expressing cells across the GC-MS metabolomic profiling (Fig. S3c,d,f,g). In a complementary LC-MS analysis, L-tyrosine and structurally related metabolites such as phenylpyruvate and benzoic acid also showed intracellular accumulation with SLC16A6^S240A^ expression (Fig. S3i,j). Consistent with the metabolomics screen, metabolic profiling by RESOLUTE in HEK-293T cells expressing WT SLC16A6 revealed significant shifts in intracellular hydroxybenzoic acid, a tyrosine derivative, and phenylalanine (Fig. S4a)^31^. In parallel, *in silico* substrate predictions by AlphaFill favored lactate and the phenylalanine derivative, hydrocinnamic acid (Fig. S4b)^32^. Our unbiased screen, along with the RESOLUTE and AlphaFill datasets, led to the hypothesis that SLC16A6 may transport L-tyrosine, the amino acid necessary for melanin production.

**Figure 3.**
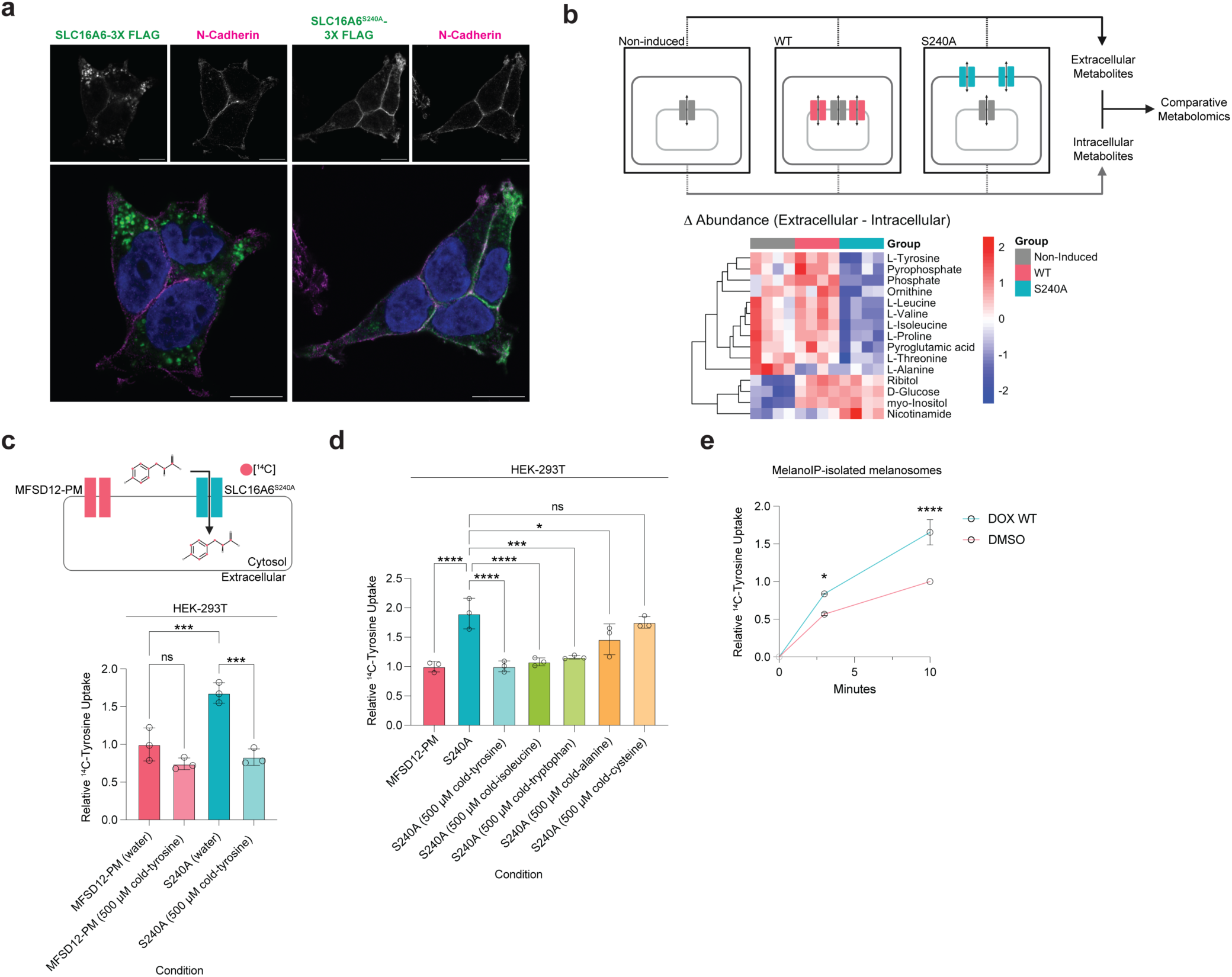
SLC16A6 is a tyrosine transporter. **(a)** SLC16A6^S240A^ reroutes SLC16A6 to the plasma membrane. Representative confocal images of dox-induced wildtype (WT) or SLC16A6^S240A^ in HEK-293T cells (images representative of n=3 biological replicates; scale bar 10 µm). **(b)** *Top*, schematic of untargeted metabolomics screen. Stable HEK-293T cells with either WT or SLC16A6^S240A^ mutant were treated with either DMSO or 500 ng mL^−1^ doxycycline and incubated for 24 h. Media was replaced and incubated for an additional four hours prior to intracellular and extracellular metabolite extractions. *Bottom*, heat map showing metabolites with altered extracellular-to-intracellular abundance differences between groups. Delta abundance was calculated as log2-transformed extracellular abundance minus log_2_-transformed intracellular abundance for each metabolite, followed by z-score normalization. **(c)** *Top*, schematic of [^14^C]-tyrosine cellular uptake assay. *Bottom*, HEK-293T cells expressing either MFSD12-PM or SLC16A6^S240A^ were incubated in a buffer containing a LAT1 inhibitor. Cold-tyrosine (5 µM for transport assay; 500 µM for competition) and 1 µCi mL^−1 14^C-labeled tyrosine were added to initiate transport. Assay was performed for 10 min prior to washes and scintillation. Two-way ANOVA was used for statistical analysis (\*\*\**P* < 0.001; n=3 biological replicates). **(d)** As in (c) but cold-isoleucine, -tryptophan, -alanine, -cysteine at 500 µM were added to the SLC16A6^S240A^-expressing cells. Two-way ANOVA was used for statistical analysis (\**P* < 0.05; \*\*\**P* < 0.001; \*\*\*\**P* < 0.0001; n=3 biological replicates). **(e)** L-tyrosine uptake in isolated melanosomes. MelanoIP-isolated melanosomes were extracted from cells treated with DMSO or doxycycline to induce WT SLC16A6 expression. Melanosomes were incubated for indicated times in a buffer containing ATP and Mg^2+^ to promote uptake, before washing and scintillation. Two-way ANOVA was used for statistical analysis. Data normalized to DMSO-treatment at 10 min (\**P* < 0.05, \*\*\*\**P* < 0.0001; n=3 biological replicates).

To examine whether SLC16A6 was sufficient for tyrosine transport, we asked whether doxycycline-inducible expression of SLC16A6 ^S240A^ increased the cellular uptake of tyrosine. To strengthen the signal-to-noise ratio of these uptake assays, we included an inhibitor of LAT1, the major neutral amino acid transporter of the plasma membrane^33,34^. HEK-293T cells expressing SLC16A6^S240A^ exhibited a clear increase in radiolabeled tyrosine uptake as compared to the control MFSD12-PM, the melanosomal cysteine transporter re-localized to the plasma membrane (Fig. 3c). Importantly, the increase in transport was blocked by the addition of excess unlabeled or “cold” tyrosine, suggesting that tyrosine transport is saturable, a cardinal feature of specific substrate transport (Fig. 3c). This increase in tyrosine uptake was also observed in comparison to non-induced SLC16A6^S240A^ HEK-293Ts as well as HEK-293Ts with induction of WT SLC16A6, which predominantly localizes to internal compartments rather than the plasma membrane (Fig. 3a, Fig. S5a,b). Moreover, SLC16A6^S240A^ enhances tyrosine uptake even in the absence of the LAT1 inhibitor (Fig. S5c), further supporting its sufficiency to transport tyrosine.

The melanosomal tyrosine transporter was previously described as a transport system capable of transporting other aromatic and large neutral amino acids^2^. To test whether SLC16A6 shares this substrate profile, we performed amino acid competition assays in cellular uptake assays. Again, cold tyrosine blocked [^14^C]L-tyrosine uptake (Fig. 3d). We found that unlabeled isoleucine and tryptophan also competed with radiolabeled tyrosine uptake, with mean percent inhibition of 91% and 83%, respectively, whereas unlabeled alanine (38%) and cysteine (11%) were less effective at blocking labeled tyrosine uptake (Fig. 3d, Fig. S5d). The pattern of amino acid competition was very similar to the reported range of inhibition for the previously described, but unidentified, melanosomal tyrosine transport system^2^. We further examined previously reported substrates of SLC16A6 and observed that excessive unlabeled taurine and β-hydroxybutyrate failed to compete with labeled tyrosine uptake and therefore are unlikely to be high-affinity substrates (Fig. S5e).

To directly test whether SLC16A6 was sufficient for tyrosine uptake into melanosomes, we induced WT SLC16A6 expression in the mouse melanoma B16F10 cell line stably expressing the MelanoIP system^8^. Upon isolation of melanosomes by MelanoIP, we performed a radiolabeled L-tyrosine uptake assay and observed increased uptake of [^14^C]L-tyrosine when compared to control melanosomes (Fig. 3e). Importantly, when excess cold tyrosine was added to the reaction, or the reaction was performed at 4°C, we did not observe this increase in tyrosine uptake (Fig. S5f). Together, these results are consistent with SLC16A6 being a melanosomal tyrosine transporter.

### SLC16A6 is required for melanosome formation

Given that melanin biosynthesis relies on tyrosine transport into early-stage melanosomes, we next assessed the phenotype of melanoma cells upon loss of *SLC16A6*. Upon *SLC16A6* knock out (Fig. S6a), human melanoma MeWo cells changed morphology (Fig. 4a) and showed significantly reduced proliferation compared with parental MeWo cells, but without any evidence of cell death (Fig. 4b, Fig. S6b). *SLC16A6* knockout MeWo and B16F10 cells were also devoid of apparent pigmentation (Fig. 4c), suggesting that SLC16A6 might be necessary for melanogenesis. Proteomic profiling of *SLC16A6* knockout cells showed systemic downregulation of most melanosomal components (Fig. 4d). Accordingly, early melanogenesis components like PMEL and MLANA/MART-1, and late melanogenesis components like TYR, TYRP1, and DCT/TYRP2 were all undetectable at the protein level in MeWo and B16F10 knockout cells (Fig. 4e, Fig. S6c).

**Figure 4.**
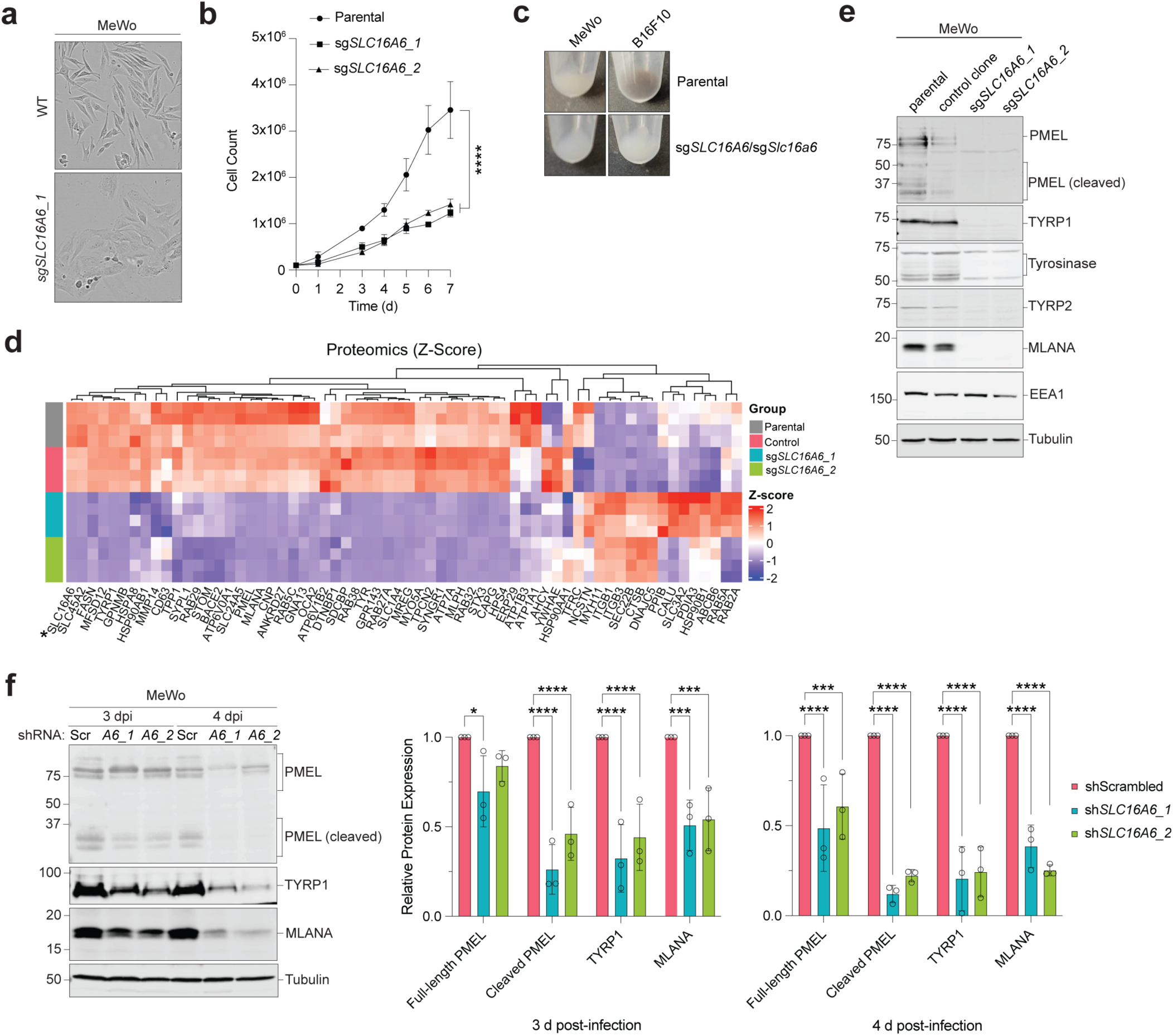
SLC16A6 is necessary for melanosomes. **(a)** Brightfield images of WT or *SLC16A6* knockout cells. **(b)** Proliferation assay of parental versus *SLC16A6* knockout clone MeWo cells. One hundred thousand cells were plated at day 0, and counted at days 1, 3, 4, 5, 6, and 7 by hemocytometer (\*\*\*\**P* < 0.0001; n=3, biological replicates). **(c)** Representative photographs of melanotic parental versus amelanotic *SLC16A6* knockout clones in human MeWo and mouse B16F10 melanoma cells. **(d)** TMT quantitative proteomic heatmap of MeWo parental, control clone, and two *SLC16A6* knockout clones. Proteins highlighted are isolated from whole cell proteomics dataset using the gene ontology GO:0042470, melanosome. Asterisk denotes SLC16A6. **(e)** Representative immunoblots of indicated melanogenesis components in parental, control clone, and two *SLC16A6* knockout MeWo clones. **(f)** *Left,* representative immunoblots of melanogenesis components in MeWo cells three and four days post-infection with indicated shRNA lentiviruses (*Scr*, scrambled shRNA control; *A6_1*, shRNA targeting SLC16A6 #1; *A6_2*, shRNA targeting SLC16A6 #2; n=3 biological replicates). *Right*, quantification of relative protein expression of shRNA treatment in at least n=3 biological replicates. Each protein is normalized to control alpha-tubulin (\**P* < 0.05; \*\*\**P* < 0.001; \*\*\*\**P* < 0.0001).

We next tested whether acute loss of *SLC16A6* resulted in a similar loss of pigmentation and melanogenesis. We observed decreases in melanosomal components at 3 days post-infection with lentiviral shRNAs targeting *SLC16A6* (Fig. 4f, Fig. S6d-f). Melanosomal biogenesis requires proper PMEL processing by BACE2 to form amyloid-like fibrils to act as both structural units for developing melanosomes and for melanin deposition^35^. Interestingly, PMEL cleavage defects were the most immediate phenotype observed following acute *SLC16A6* knockdown, followed by the sequential loss of MLANA and TYRP1 levels (Fig. 4f, Fig. S6d). These results are consistent with SLC16A6 functioning in early melanogenesis, where the lack of tyrosine transport triggers a cascade leading to melanogenesis defects.

## Conclusion

Our findings establish the orphan transporter SLC16A6 as a long-sought melanosomal tyrosine transporter, answering a biochemical question first proposed in 1996^2^. We determined that SLC16A6 acts as an early melanogenesis solute carrier whose expression is driven by the melanocyte lineage-defining transcription factors SOX10 and MITF. Importantly, we show SLC16A6 is sufficient to transport L-tyrosine in both whole-cell uptake assays using the plasma membrane-rerouted SLC16A6^S240A^ mutant and melanosome-specific transport assays using WT SLC16A6 expression. Furthermore, our SLC16A6^S240A^ assays robustly mimic the competitive inhibition profile for the reported melanosomal tyrosine transporter: L-tyrosine uptake is competed by L-tyrosine, L-isoleucine and L-tryptophan, while being less sensitive to L-alanine and L-cysteine. Ultimately, this alignment between data presented 30 years apart provides strong support for SLC16A6 being the protein that was previously studied as the melanosomal tyrosine transporter.

Unexpectedly, our data reveal a dependency between tyrosine compartmentalization and organelle biogenesis, in which the loss of *SLC16A6* triggers an almost complete loss of core melanosomal components. Foremost among these affected components is PMEL. PMEL is essential for melanosomal structure and function, as cleaved PMEL forms amyloid fibril-like scaffolds that serve both structural roles and sites for melanin deposition^35,36^. Upon *SLC16A6* knockdown, PMEL cleavage becomes impaired, resulting in accumulation of uncleaved PMEL and subsequent loss of total PMEL protein. Yet, unlike *PMEL* deletions^37^, *SLC16A6* knockdowns and knockouts deplete all other melanosomal components as well. While intriguing and consistent with its role in melanin synthesis, this phenotype makes it impossible for us to define the effects of loss of *SLC16A6* function in mature melanocytes, as they are lost concomitant with the loss of *SLC16A6* expression.

Our data suggest that the loss of tyrosine transport culminates in melanosomal failure, directly linking organelle homeostasis to the synthesis of its central metabolite. While mutations or deletions in lysosomal transporters such as *CTNS* or *NPC1* cause cystine crystallization and cholesterol accumulation, respectively, these organelles remain intact, albeit dysfunctional^38–40^. Similarly, the loss of mitochondrial mitoferrins, SLC25A28 and SLC25A37, profoundly impairs oxidative respiration prior to cell death but does not abolish mitochondrial biogenesis^41,42^. In contrast, lysosome related organelles appear uniquely sensitive to their internal environment. This phenomenon is observed with Weibel-Palade bodies, where specific mutations (e.g., A1716P and C2190Y) in von Willebrand factor (*vWF*) disrupt the biogenesis of these organelles^43^. Similarly, in type 2 alveolar cells, deletion of the dipalmitoylphosphatidylcholine (DPPC) transporter *Abca3* halts the formation of lamellar bodies^44,45^. Whether other intracellular metabolite transporters regulate organelle identity in a manner similar to SLC16A6 and ABCA3 remains an open question. Ultimately, identifying SLC16A6 as a key regulator of melanogenesis provides a new molecular framework for understanding human pigmentation disorders and offers a compelling new target for melanocyte-driven pathologies.

## Methods

### Cell Culture

HEK-293T and SK-MEL-30 cells were from ATCC. MeWo cells were a gift from Dr. Martin McMahon. B16F10 cells were a gift from Dr. Minna Roh-Johnson. HEK-293Ts and B16F10 cells were cultured in DMEM with L-Glutamine, 4.5g L^−1^ glucose, and sodium pyruvate (Gibco) supplemented with 10% FBS (Sigma). MeWo and Sk-Mel-30 cells were cultured in RPMI 1640 (Gibco) supplemented with 10% FBS. Dox-inducible HEK-293T and B16F10 cells were cultured in DMEM (Gibco) supplemented with 10% tetracycline-free FBS (Takara). All cells were propagated at 37°C with 5% CO2. Cells were routinely tested for mycoplasma contamination.

### Virus production and generation of stable cell lines

Lentiviruses were produced by co-transfection of HEK-293T cells in a 10 cm plate with a target lentiviral vector (5 μg), pMDG.2 (1.67 μg), pMDLg/pRRE (1.67 μg), and pRSV-REV (1.67 μg) plasmids. Virus containing supernatants were collected 48 and 72 h post-transfection, spun at 1000 *g* for 5 min, and then filtered through a 0.45 μm filter. For infection, approximately 10^6^ cells were incubated with 1-5 mL viral supernatant, 10 μg mL^−1^ polybrene and culture medium. Viral media was removed after 24 h and replaced with fresh culture medium for 24 h, prior to the addition of a selection agent. Selection was maintained for a minimum of 3 d or until all non-infected control cells had undergone apoptosis.

### Knockout cell line generation

The following plasmid was acquired from Addgene:

pSpCas9(BB)-2A-GFP (PX458) was a gift from Feng Zhang (Addgene plasmid #48138 ; http://n2t.net/addgene:48138 ; RRID:Addgene_48138)^46^

The following sgRNA sequences were subcloned into the above construct:

SLC16A6_sgRNA1_exon1_sense: CACCGAGGCACTTCAGTATACACAT
SLC16A6_sgRNA1_exon1_antisense: AAACATGTGTATACTGAAGTGCCTC
SLC16A6_sgRNA2_exon1_sense: CACCGAAATGTCTTGATGATGCCGT
SLC16A6_sgRNA2_exon1_antisense: AAACACGGCATCATCAAGACATTTC
mSlc16a6_sgRNA1_exon1_FWD: CACCGGCGACCCGCGATGACAGCG
mSlc16a6_sgRNA1_exon1_REV: AAACCGCTGTCATCGCGGGTCGCC
mSlc16a6_sgRNA2_exon1_FWD: CACCGCACCAGCCGATGTCCGAAG
mSlc16a6_sgRNA2_exon1_REV: AAACCTTCGGACATCGGCTGGTGC

PX458 constructs were transiently transfected into MeWo or B16F10 cells using Lipofectamine 3000 (Invitrogen) according to the manufacturer’s protocol. Cells were GFP flow sorted single cell into 96-well plates and outgrown clones were screened for *SLC16A6* knockout.

### cDNA constructs

The following plasmids were acquired from Addgene:

1. pEGFP-N1-MITF-M was a gift from Shawn Ferguson (Addgene plasmid #38131 ; http://n2t.net/addgene:38131 ; RRID: Addgene_38131)^47^
2. pEGFP-N1-MITF-A was a gift from Shawn Ferguson (Addgene plasmid #38132 ; http://n2t.net/addgene:38132 ; RRID: Addgene_38132)^47^
3. pEGFP-N1-MITF-D was a gift from Shawn Ferguson (Addgene plasmid #38133 ; http://n2t.net/addgene:38133 ; RRID: Addgene_38133)^47^

Wildtype (WT) SLC16A6-3X FLAG was synthesized by Twist Biosciences using the mRNA sequence for *Homo sapiens* solute carrier family 16 member 6 (SLC16A6), transcript variant 1 (NCBI Accession: NM_001174166). The S240A mutant was subsequently generated from the WT construct via site-directed mutagenesis.

### shRNA-mediated knockdown

shRNA in a pLKO.1 backbone targeting *SLC16A6* was purchased from Sigma. The following sequences were used:

shSLC16A6_1: CTACGGCATCATCAAGACATT
shSLC16A6_2: GTAACCATCCTATCACAATAT

Knockdowns were confirmed by quantitative PCR (qPCR). RNA was isolated from individual 6 cm plates that were 80% confluent using Trizol and an RNA Extraction Kit (Zymo). cDNA was generated using a High-Capacity cDNA Reverse Transcription Kit (Applied Biosystems) and diluted to 10 ng μL^−1^ before using 2 μL for RT-qPCR reaction. *SLC16A6*, *PolR2a*, and *RPLP0* transcript levels were assayed with SYBR green qPCR mix (Roche), using the following primer sets:

qPCR_SLC16A6_F: GGCATCATCTCTGGTCTGGG
qPCR_SLC16A6_R: TGGTGCGAAAGCAAACACAG
qPCR_PolR2a_F: AACACGGGCAGAGATCCAAG
qPCR_PolR2a_R: TTCATGACTTCACCCCGCTC
qPCR_RPLP0_F: CTGTTGGCCAATAAGGTGCC
qPCR_RPLP0_R: GGAGGTCTTCTCGGGTCCTA

### Immunoblots

MeWo and SK-Mel-30 cells were washed and scraped in PBS. Cells were spun at 1000 *g* for 3 min and supernatant discarded. Pellets were lysed in 1% Triton X-100 in 50mM Tris-HCl pH 7.4, 150mM NaCl, supplemented with mammalian protease inhibitor cocktail (mPIC) at 1:100 (Sigma-Aldrich). For tyrosinase immunoblots, cells were lysed in 0.1% n-Dodecyl-β-D-maltoside (Anatrace) instead of 1% Triton X-100. All cells were lysed on ice for 30 min before centrifuging at 16100 *g* for 10 min. Protein quantifications were analyzed by Pierce BCA Protein Assays (ThermoFisher Scientific). 20-50 μg of protein was loaded per well on SDS-PAGE, transferred to 0.45 μm nitrocellulose by wet transfer, blocked in 5% non-fat milk. All primary antibodies were incubated overnight at 4 °C, at a common ratio of 1:2000 except for alpha-tubulin (1:10000). Secondary antibodies Alexafluor-conjugated goat anti-rabbit (IRDye800, Rockland) and/or goat anti-mouse (Alexafluor-680, Life Technologies) antibodies were incubated for 1 hour at room temperature. Immunoblots were imaged using a LI-COR Odyssey CLx imager.

### Immunofluorescence

MeWo and HEK-293T cells were plated on 18×18 #1 coverslips and 18×18 #1, poly-l-lysine-coated coverslips, respectively. Cells were washed twice in PBS before fixing with 4% paraformaldehyde for 15 min. Coverslips were washed three times before permeabilization with 100% ice-cold methanol at −20 °C for 5 min. Cells were washed and blocked with 3% Fatty-Acid Free Bovine Serum Albumin for at least one hour at 4 °C. The following primary antibodies were incubated overnight at the indicated ratios: mouse anti-FLAG M2 (1:1000; Sigma), rabbit anti-FLAG (1:500; Cell Signaling Technologies), mouse anti-EEA1 (1:500; Cell Signaling Technologies), goat anti-LAMP2 (1:500; Biotechne), rabbit anti-SLC16A6 (1:250; Sigma), rabbit anti-N-Cadherin (1:500; Cell Signaling Technologies), and mouse anti-TRP1 (1:1000; Abcam). Donkey anti-rabbit IgG (H+L) highly cross-adsorbed Alexafluor 488 (1:500), Donkey anti-mouse IgG (H+L) highly cross-adsorbed Alexafluor 568 (1:500), and Donkey anti-goat IgG (H+L) cross-adsorbed (1:500) secondary antibodies were incubated for 1 hour at room temperature and washed three times with PBS. Coverslips were staged with ProLong Diamond Antifade with DAPI (Fisher Scientific). Images were captured on a Zeiss LSM 880 confocal laser scanning microscope. Line scan analyses were performed in Fiji. Whole cell pearson correlation coefficients were analyzed in Fiji by the Coloc 2 plugin. Line scan pearson correlation coefficients were calculated using GraphPad Prism 10.

### Cancer Dependency Map analyses

Correlation analysis of SLC16A6 mRNA and protein expression were performed on Custom Analyses on the DepMap portal. For mRNA correlation, the Expression (Short-read) Public 26Q1 was used to identify transcripts with high correlations to *SLC49A3* and *SLC16A6* transcript abundance. The Harmonized MS CCLE Gygi dataset was used to identify proteins with high correlations to SLC16A6 protein abundance. Lineage expression analysis of *SLC16A6* was performed by downloading the expression (short-read) public 26Q1 dataset.

### TRoUT-FISH screen

The following CRISPR library was acquired from Addgene:

1. Human Epigenetic Knockout Library was a gift from Kivanc Birsoy (Addgene #162256 ; http://n2t.net/addgene:162256 ; RRID:Addgene_162256)^48^

Lentiviruses were produced in HEK-293T cells by transfecting plasmids and packaging plasmids (psPAX2 and VSVG) with polyethylenimine transfection reagent. Media was collected every 24 h for 96 h total. Media was pooled, filtered with 0.45 µm PES filters, aliquoted, and frozen at −80 °C.

The titer of lentivirus supernatant was determined by infecting target cells at various amounts of virus with polybrene followed by spinfection (30 °C for 60 min at 1200 *g*). The next day cells were reseeded and treated with puromycin. Viability was assessed 3 d post infection. Cells were infected at a Multiplicity of Infection (MOI) of ∼0.4 via spinfection, and treated with puromycin 24 h post infection and puromycin selection persisted for 7 d. Cells were given fresh media for 6 h followed by fixation with cold 4% PFA.

Cells were permeabilized with PBS containing 0.1% Triton X-100 and Hoechst 33342 for 10 min rocking at 4 °C. Cells were incubated with 43 °C wHyb buffer ^49,50^ for 30 min, then incubated overnight at 43 °C in hybridization buffer containing SABER-FISH PER purified DNA oligonucleotides. Subsequently, cells were washed two times with 43 °C wHyb buffer, one time with 2X SSC buffer, and incubated at 30 °C for 1 h in branch hybridization buffer containing SABER-FISH PER purified branch DNA oligonucleotides. After branching, cells were washed with 2X SSC and 37 °C PBS before incubation for 10 min in PBS containing fluorescently labeled secondary DNA oligonucleotides. After staining, cells were washed two times with 37 °C PBS and resuspended in 0.1% BSA (Millipore, Catalog: 126609-5GM) in PBS for flow cytometry.

Events were gated based on FSC and SSC to remove debris, then gated on DNA dye (violet laser) to ensure only singlets were sorted. A reference gene (EEF2) was used as a control to ensure cell quality and adequate staining. Cells were sorted based on fractions of the target gene distribution, sorting the lowest 10% and highest 10% of the total population. An unsorted population was also acquired. After sorting cells were incubated in genomic DNA isolation buffer (0.5% SDS, 10 mM TRIS pH 7.5, 10 mg mL^−1^ Proteinase K) and incubated at 56 °C for ≥ 36 h with gentle agitation. Genomic isolation was performed with the Qiagen DNA Mini Kit (Catalog: 51304). Libraries were generated by PCR amplification and sequenced by 150 bp paired-end sequencing at the University of Utah High-Throughput Genomics Core Facility. Sequencing reads were quantified using BBtools Seal and analysis was performed by MAGeCK^51,52^.

### ChIP-Seq and ChIP-Atlas analysis

MITF ChIP sequencing data from MeWo KRAB cells^25^ was extracted and analyzed on Integrated Genome Viewer (2.18.2). MITF binding at gene loci was assessed using ChIP-Atlas^53–55^. We queried “Peak Browser”, restricting the search to the track class “TFs and others”, cell type class “Epidermis”, track type “MITF”, and cell type “501A” using the hg38 genome assembly. Peaks were called with a MACS2 Q-value threshold of 10^−50^ (ChIP-Atlas significance score of 50). MITF ChIP-Seq data from GSE172383 (SRX10640872 and SRX10640873) was queried within 5-10 kb of a transcriptional start site of *SLC16A6*, *TYR*, *SLC45A2*, and *DCT*. Genome tracks images were generated on Integrated Genome Viewer (2.18.2).

### LC-MS Metabolomics

Cell pellets were extracted in 200 µL of 70% ACN, 5% MeOH, and 25% H2O + 0.1% NH_4_OH, 0.1 ug mL^−1^ D9-carnitine, 2.5 ul of labeled amino acid mix (Cambridge Isotope Laboratories, MSK-A2-1.2), and 1 ug mL^−1^ D4-succinate. Samples were sonicated on ice for 5 min, vortexed, and centrifuged at 20,000 *g* for 10 min at 4 °C. Supernatants were collected and stored at −80 °C prior to analysis/injection. Pooled quality control samples were prepared by combining equal volumes of each sample and process blanks were made using only extraction solvent and standards. Quantification of metabolites was performed by LC-MS using a SCIEX 7600 Zeno-ToF coupled to an Agilent 1290 Infinity II HPLC system. Chromatographic separation was achieved on a Waters BEH zhilic 2.1 × 100 mm column with Phenomenex Krudkatcher Ultra with a binary mobile phase of 10mM ammonium carbonate in water (A) and acetonitrile/water (95:5, v/v) (B) with using an optimized X-min gradient (min/%B: 1/99, 2/85, 5/75, 8/60, 10/40, 11/1, 12/1). Metabolites (n=390) were identified and confirmed by both high-resolution multiple reaction monitoring and retention time of analytical standards. Chromatogram integration and quantification were performed using SCIEX MultiQuant software.

### GC-MS Metabolomics

Cell pellets and media samples were extracted in cold 90% MeOH containing 0.67 ug mL^−1^ D4 succinic acid and 20 ug mL^−1^ D27 myristic acid, incubated at −20 °C for 1 h, and centrifuged at 20,000 *g* for 10 min at 4 °C. Supernatants were collected and dried en vacuo. Pooled quality control samples and process blanks were prepared as above. Dried samples were suspended in O-methoxylamine hydrochloride (MOX) (MP Bio #155405) in dry pyridine (EMD Millipore #PX2012-7) and incubated for one hour at 37 °C before subsequent derivatization with N-methyl-N-trimethylsilyltrifluoracetamide (MSTFA with 1% TMCS, Thermo #TS48913) for 30 minutes at 37 °C. After incubation, samples (1 µL) were injected in the split mode (5:1 split ratio for most metabolites) at an inlet temperature of 250 °C. Samples were analyzed on an Agilent 5977b GC-MS MSD-HES system fit with an Agilent 7693A automatic liquid sampler. Chromatographic separation was performed with a DB-5MS capillary column (30 m × 0.25 mm × 0.25 µm) with a 10 m Duraguard precolumn (Agilent Technologies) using helium carrier gas (1 mL/min) and a temperature program of 60 °C (1 min hold), ramping at 10 °C min^−1^ (10 min hold) to 325 °C (10 min hold). Metabolites (n=83) were identified and quantified by retention index and characteristic fragment ion matching against an in-house pure standard library, the Fiehn library, and the NIST library. Data was collected using MassHunter software (Agilent).

### Whole-Cell Proteomics

#### Sample Preparation for Mass Spectrometry

Samples for protein analysis were prepared essentially as previously described^56,57^. Proteomes were extracted using a buffer containing 200 mM EPPS pH 8.5, 8M urea, 0.1% SDS and protease inhibitors. Following lysis, 25 µg of each proteome was reduced with 5 mM TCEP. Cysteine residues were alkylated using 10 mM iodoacetimide for 20 min at RT in the dark. Excess iodoacetimide was quenched with 10 mM DTT. A buffer exchange was carried out using a modified SP3 protocol^58^. Briefly, ∼250 µg of Cytiva SpeedBead Magnetic Carboxylate Modified Particles (65152105050250 and 4515210505250), mixed at a 1:1 ratio, were added to each sample. 100% ethanol was added to each sample to achieve a final ethanol concentration of at least 50%. Samples were incubated with gentle shaking for 15 min. Samples were washed three times with 80% ethanol. Protein was eluted from SP3 beads using 200 mM EPPS pH 8.5 containing Lys-C (Wako, 129-02541). Samples were digested overnight at room temperature with vigorous shaking. The next morning trypsin (ThermoFisher Scientific) was added to each sample and further incubated for 6 h at 37 °C. Following digestion, an equal volume of each sample was pooled to generate a pooled sample to be used as a bridge between two TMTPro 16-plex experiments. Acetonitrile was added to each sample to achieve a final concentration of ∼33%. Each sample was labelled, in the presence of SP3 beads, with ∼62.5 µg of TMTPro reagents (ThermoFisher Scientific). Following confirmation of satisfactory labelling (>97%), excess TMT was quenched by addition of hydroxylamine to a final concentration of 0.3%. The full volume from each sample was pooled and acetonitrile was removed by vacuum centrifugation for 1 hour. The pooled sample was acidified and peptides were de-salted using a Sep-Pak 50mg tC18 cartridge (Waters). Peptides were eluted in 70% acetonitrile, 1% formic acid and dried by vacuum centrifugation.

#### Basic pH reversed-phase separation (BPRP)

TMT labeled peptides were solubilized in 5% ACN/10 mM ammonium bicarbonate, pH 8.0 and ∼300 µg of TMT labeled peptides were separated by an Agilent 300 Extend C18 column (3.5 μm particles, 4.6 mm ID and 250 mm in length). An Agilent 1260 binary pump coupled with a photodiode array (PDA) detector (Thermo Scientific) was used to separate the peptides. A 45 min linear gradient from 10% to 40% acetonitrile in 10 mM ammonium bicarbonate pH 8.0 (flow rate of 0.6 mL min^−1^) separated the peptide mixtures into a total of 96 fractions (36 seconds). A total of 96 Fractions were consolidated into 24 samples in a checkerboard fashion and vacuum dried to completion. Each sample was desalted via Stage Tips and re-dissolved in 5% FA/ 5% ACN for LC-MS3 analysis.

#### Liquid chromatography separation and tandem mass spectrometry (LC-MS3)

Proteome data were collected on an Orbitrap Fusion Lumos mass spectrometer (ThermoFisher Scientific) coupled to a Proxeon EASY-nLC 1000 LC pump (ThermoFisher Scientific). Fractionated peptides were separated using a 120 min gradient at 550 nL min^−1^ on a 35 cm column (i.d. 100 μm, Accucore, 2.6 μm, 150 Å) packed in-house. MS1 data were collected in the Orbitrap (60,000 resolution; maximum injection time 50 ms; AGC 4 × 10^5^). Charge states between 2 and 6 were required for MS2 analysis, and a 120 s dynamic exclusion window was used. Top 10 MS2 scans were performed in the ion trap with CID fragmentation (isolation window 0.5 Da; Rapid; NCE 35%; maximum injection time 50 ms; AGC 2 × 10^4^). An on-line real-time search algorithm (Orbiter) was used to trigger MS3 scans for quantification^59^. MS3 scans were collected in the Orbitrap using a resolution of 50,000, NCE of 55%, maximum injection time of 200 ms, and AGC of 3.0 × 10^5^. The close out was set at two peptides per protein per fraction^59^.

#### Data analysis

Raw files were converted to mzXML, and monoisotopic peaks were re-assigned using Monocle^60^. Searches were performed using the Comet search algorithm against a mouse database downloaded from Uniprot in May 2021. We used a 50 ppm precursor ion tolerance, 1.0005 fragment ion tolerance, and 0.4 fragment bin offset for MS2 scans collected in the ion trap. TMTpro on lysine residues and peptide N-termini (+304.2071 Da) and carbamidomethylation of cysteine residues (+57.0215 Da) were set as static modifications, while oxidation of methionine residues (+15.9949 Da) was set as a variable modification.

Each run was filtered separately to 1% False Discovery Rate (FDR) on the peptide-spectrum match (PSM) level. Then proteins were filtered to the target 1% FDR level across the entire combined data set. For reporter ion quantification, a 0.003 Da window around the theoretical m/z of each reporter ion was scanned, and the most intense m/z was used. Reporter ion intensities were adjusted to correct for isotopic impurities of the different TMTpro reagents according to manufacturer specifications. Peptides were filtered to include only those with a summed signal-to-noise (SN) ≥ 160 across all TMT channels. The signal-to-noise (S/N) measurements of peptides assigned to each protein were summed (for a given protein). These values were normalized so that the sum of the signal for all proteins in each channel was equivalent thereby accounting for equal protein loading.

### MelanoIP Proteomics

Ten million MeWo cells transduced with HA-MelanoIP or myc-MelanoIP were plated in 15 cm dishes and expanded for 2 d in the presence of 10 µM forskolin. Cells were harvested by scraping in ice cold PBS, pelleted by centrifugation at 1000 g for 3 min at 4 °C, and resuspended in 200 µL PBS supplemented with protease inhibitors (Roche). Cells were then homogenized with a handheld rotary pestle (Kimble-Chase) for 45 seconds before adjusting the total volume to 1 mL with PBS and protease inhibitors and spinning at 1000 *g* for 5 min at 4 °C to clarify. Supernatants were transferred into new 1.5 mL tubes containing 100 µL of PBS-washed anti-HA magnetic beads (Themo Fisher) and incubated rocking for 10 minutes at 4 °C. Beads were washed 3 times with cold PBS with the final wash transferred to a new 1.5 mL tubes. Beads were then eluted with 5% SDS, 50 mM HEPES, pH 7.4, 50 mM NaCl for 10 minutes at 4 °C.

#### Proteomic sample preparation

Melano-IP samples (3 independent control Myc and experimental HA enrichments) were reduced with 1 mM dithiothreitol at 56 °C for 10 min and alkylated with 2.25 mM iodoacetamide at room temperature for 20 min. Reduced and alkylated proteins were precipitated onto S-Trap micro columns (ProtiFi) following the manufacturer’s instructions. Purified proteins were digested with 2 µg of a 1:5 w/w Trypsin:LysC mixture (Worthington and Sigma, respectively) at 37 °C for 4 h. Peptides were eluted from the columns following the manufacturer’s instructions. Fourteen percent of each peptide sample was loaded onto Evotip Pure (EvoSep) following the manufacturer’s instructions.

#### LC-MS analysis

Peptides were loaded onto a 15 cm x 150 µm x 1.9 µm column (Evosep) and separated across a 44-min gradient (30SPD) using the Evosep One chromatography system. Peptides were introduced into a timsTOF HT mass spectrometer (Bruker) via a Captive Spray 2 ion source using a 20 µm PepSep emitter. The mass spectrometer was operated in Data Independent Acquisition (DIA) mode using a custom dia-PASEF method (50 ms ramp time, 1 MS1 ramp, 20 MS/MS ramps, 60 MS/MS windows, (350 -1250) m/z mass range, (0.75 – 1.3) 1/k_0_ mobility range).

#### Database search and protein quantification

Raw DIA files for all 6 samples were used to build a custom spectral library in FragPipe (version 24.0). Pseudo-MS/MS spectra were extracted using diaTracer (2.2.1)^61^ and searched using the MSFragger algorithm (4.4.1)^62^ against a human database downloaded from UniProt on 5/22/2026. The search parameters included: 20 ppm precursor and fragment ion tolerances and up to 2 missed cleavages using a strict trypsin cleavage definition. Carbamidomethylation of cysteine (+57.0215 Da) was set as static modification. N-terminal protein acetylation (+42.0106 Da), oxidation of methionine (+15.9949 Da), pyro-glutamination of glutamine or loss of ammonia of cysteine at the peptide N-terminus (−17.0265 Da) and pyro-glutamination of glutamic acid at the peptide N-terminus (−18.0106) were set as variable modifications. Default parameters for MSBooster (1.4.14)^63^, Percolator (3.7.1)^64^, ProteinProphet^65^, and Philosopher (5.1.3-RC9)^66^ were used for protein inference and to improve peptide identification. Peptide-spectrum match (PSM) and proteins were filtered to enforce a 1% false discovery rate (FDR).

Precursors were quantified using DIA-NN (2.2.0)^67^ using the spectral library generated above, with match between runs (MBR) enabled. Precursors were filtered to enforce a 1% FDR. Relative protein abundances were estimated from precursor quantities using the R package iq (2.0.1), excluding non-proteotypic peptides. Protein abundances were normalized using the rlr method as implemented in the R package ProteoDA (2.0.0)^68^. P-values were calculated using the R limma package (3.64.3) and adjusted for multiple hypotheses using the Benjamini-Hochberg method.

### Whole-cell ^14^C-tyrosine uptake assays

Three million HEK-293T dox-inducible cells were plated in 10-cm dishes. A day later, cells were treated with DMSO or 500 ng mL^−1^ doxycycline. 48 h after, media was replaced with fresh media and incubated for 2 h. Cells were harvested by pipetting off the plate with KPBS. Cells were pelleted by centrifugation at 1000 *g* for 2 min, washed again with KPBS, and spun again. Cells were resuspended in KPBS supplemented with 125 mM sucrose and 10 µM JPH-203, a LAT1-specific inhibitor. Assays were started by the addition of cold tyrosine (5 µM for transport conditions and 500 µM for competition) and 1 µCi mL^−1^ [^14^C]tyrosine. For amino acid and reported substrate competition assays, 500 µM of cold taurine, β-hydroxybutyrate, isoleucine, tryptophan, alanine, and cysteine were added with 5 µM cold-tyrosine for transport initiation. After 10 min of incubation at 37 °C, cells were spun at 1000 *g* for 30 s at 4 °C and washed twice in ice-cold KPBS. Cell pellets were resuspended in minimal water before mixing into scintillation fluid and measuring scintillation counts (LS 6000IC, Beckman).

### MelanoIP-isolated melanosome uptake assays

Three million doxycycline-inducible B16F10 cells were plated in 15-cm dishes; two dishes used per immunoprecipitation. 24 h later, cells were treated with 10 µM forskolin and either DMSO or 500 ng mL^−1^ doxycycline. 48 h later, cell media was replaced and cells were incubated for an additional 2 h. Cells were washed twice in KPBS, and harvested, pelleted at 1000 *g* for 3 min. Cells were resuspended in 500 µL KPBS supplemented with mPIC. Cell suspension was added to a 2 mL dounce homogenizer and dounced for 40 strokes on ice. 500 µL KPBS supplemented with mPIC was added to the homogenate prior to being spun at 3000 *g* for 2 min. Supernatant protein was quantified and equal protein amounts were incubated with Pierce Magenetic HA beads (ThermoFisher) in KPBS supplemented with mPIC added to equalize volume to 1 mL. Immunoprecipitations were performed for 4 min at 4 °C with rotation before washed three times with KPBS with mPIC. Immunoprecipitated melanosomes were resuspended in ice-cold 950 µL KPBS, 125 mM Sucrose, 2 mM ATP, and 2 mM MgCl2. 5 µM cold-tyrosine and 3 µCi mL^−1^ [^14^C]tyrosine was added to start the uptake assay. Melanosomes were incubated at 37 °C for indicated times, washed three times with ice-cold KPBS before resuspension in scintillation fluid and measuring scintillation counts (LS 6000IC, Beckman). For indicated experiments, 500 µM cold-tyrosine were added, or the experiment was incubated at 4 °C and normalized to samples where 1% Triton X-100 was added.

### Cell Death Assays

5000 cells were cultured in 96-well plates (Falcon) in medium containing the cell death stain YOYO-1 Iodide (Invitrogen) diluted 1:2000 (v/v). Starting 24 h after cell seeding, phase confluency and cell death (YOYO-1 signal) were tracked in a time course at 10X magnitude using IncuCyte S3 System (Sartorius). Images were analyzed with the IncuCyte S3 Software (v2025B). Cell death quantifications were calculated as the percentage of green (YOYO-1+) area by phase area. Experiments were repeated as independent biological replicates.

### Statistical Analysis

Quantitative data were prepared in GraphPad Prism 10. Statistical comparisons were via two-tailed unpaired *t*-tests, one-way and two-way ANOVAs, all performed in Prism. All quantitative measurements are from at least n=3 biological replicates. Immunoblots and immunofluorescence are representative of experiments performed at least three independent times. Analysis of metabolomics data was performed using MetaboAnalystR^69^.

## Data availability

The data supporting the findings of this study will be made available upon publication.

## Acknowledgements

We thank members of the Rutter lab for their helpful feedback and insights. We thank members of the University of Utah Flow Cytometry Core Facility for their assistance. Metabolomics analysis was performed at the Metabolomics Core Facility at the University of Utah. Mass spectrometry equipment was obtained through NCRR Shared Instrumentation Grant 1S10OD016232-01, 1S10OD018210-01A1 and 1S10OD021505-01. This work was supported by National Institute of General Medical Sciences F32GM140525 to CNC and R35GM131854 to JR, National Cancer Institute K00CA212445 to AJB, National Institute of Arthritis and Musculoskeletal and Skin Diseases K99AR084551 and Damon Runyon Cancer Research Foundation DRG-2452-22 to CHA, and the Susan Cooper Jones Endowed Fellowship in Cancer Research to CNC. We acknowledge indirect support from Huntsman Cancer Institute’s Cancer Center Support Grant. JR is an investigator of the Howard Hughes Medical Institute.

## Contributions

Conceptualization: CNC, JR

Formal Analysis: CNC, AJB, CHA, MS, KH, JGV, AJNP, JACV, GA, NMK

Funding Acquisition: CNC, JR

Investigation: CNC, AJB, CHA, MS, KH, JGV, AJNP, JACV

Methodology: CNC, AJB, CHA, KH, JGV, GA

Project Administration: CNC, JR

Resources: SPG, DMS, JR

Software: KH, AJB

Supervision: CNC, JR

Validation: CNC, AJB, CHA, KH, JGV, AJNP

Visualization: CNC, KH

Writing – Original Draft: CNC, JR

Writing – Review and Editing: CNC, JR, AJB, CHA, MS, KH, JGV, AJNP, JACV, GA, NMK, DMS

---

**Figure S1.**
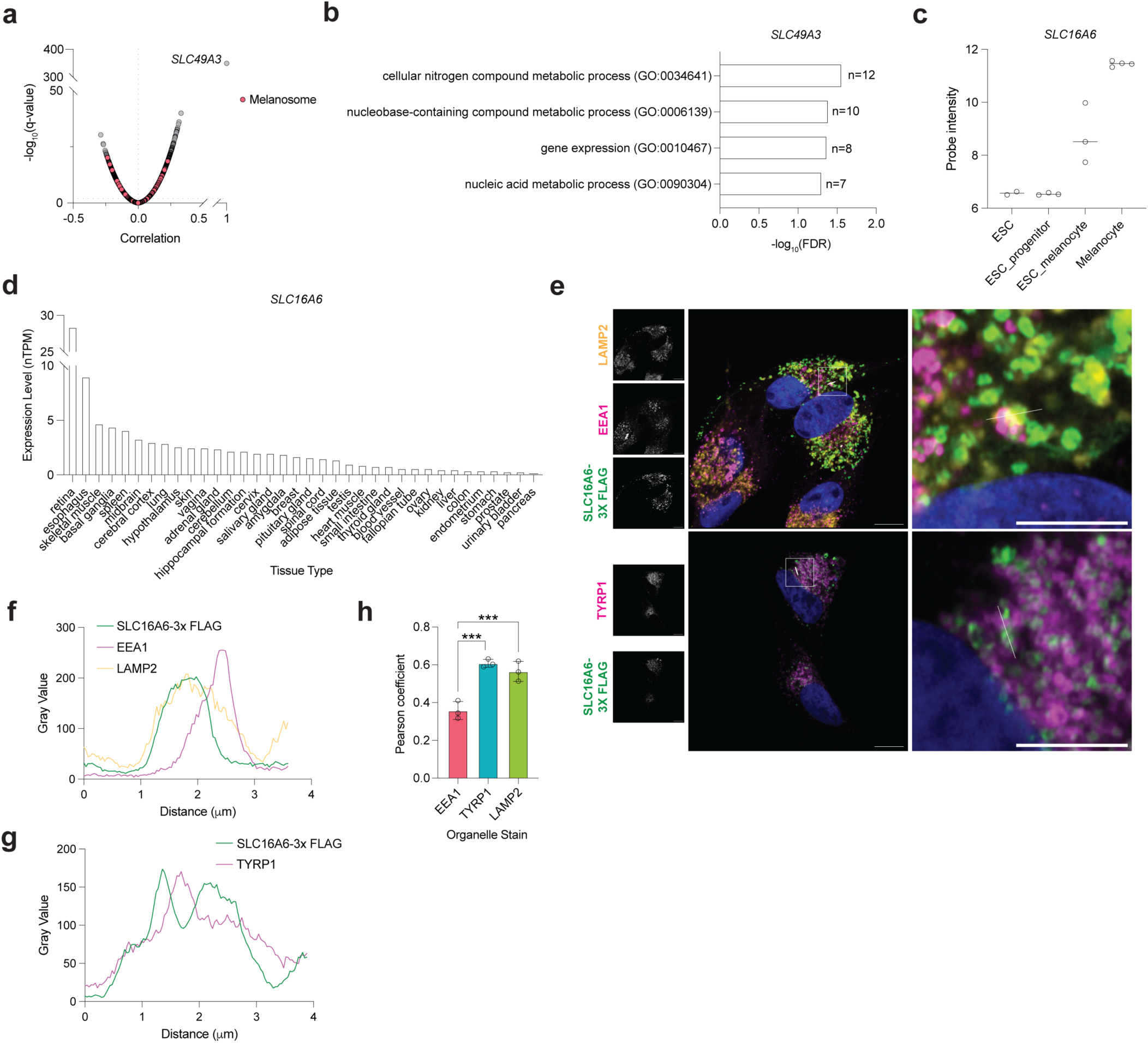
SLC16A6 is highly expressed in melanocytes and localizes to melanosomes. **(a)** *SLC49A3* transcriptome-wide correlation. Co-expression analysis of *SLC49A3* transcripts versus all other transcripts using the DepMap short-read expression dataset (26Q1). **(b)** Panther GO-slim biological process analysis of top 500 most positively correlating transcripts from (a). (**c)** *SLC16A6* expression across embryonic stem cells (ESCs), ESC-derived melanocyte progenitors, ESC-derived mature melanocytes, and primary melanocytes. Data extracted from GSE45226^70^. **(d)** *SLC16A6* expression, plotted as normalized transcript expression values, extracted from Genotype-tissue expression database. **(e)** Representative confocal microscopy images of ectopic SLC16A6-3X-FLAG colocalization (n=3 biological replicates; scale bar 10 µm). **(f)** Representative line scan analysis of SLC16A6-3X FLAG colocalization with EEA1 and LAMP2; line is indicated in (e), top image. **(g)** Same as (f) but colocalization with TYRP1; line is indicated in (e), bottom image. **(h)** Average whole cell Pearson correlation coefficients of ectopic SLC16A6-3X FLAG with indicated organelle stains (30 cells counted per trial, \*\*\**P* < 0.001, n=3 biological replicates).

**Figure S2.**
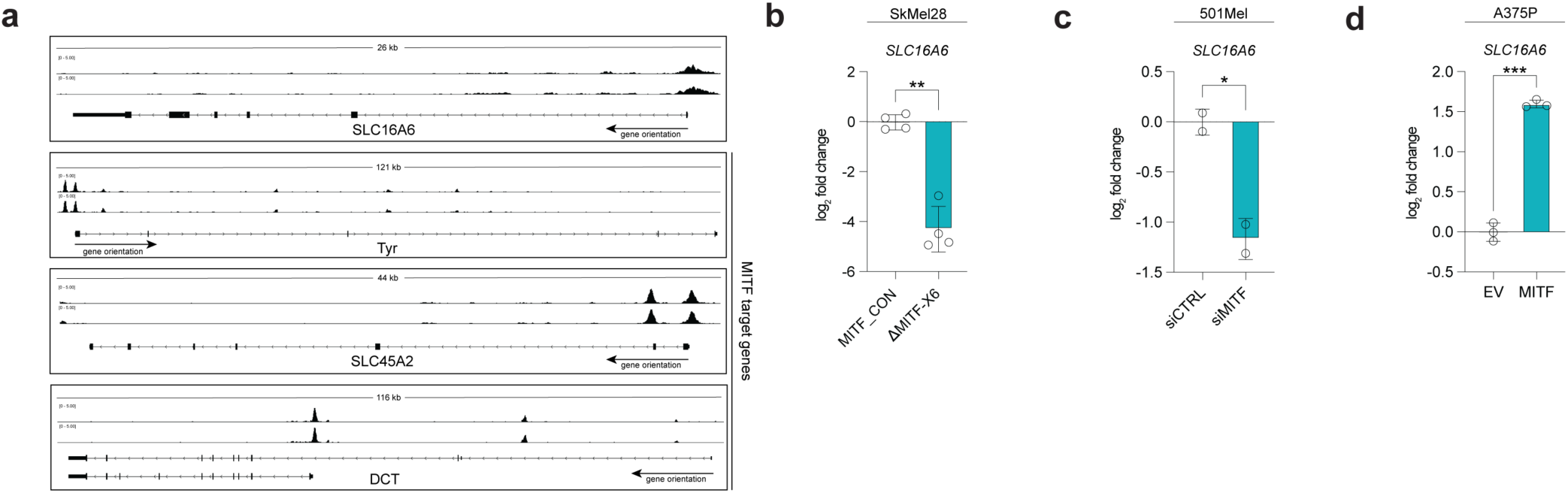
MITF binds the promoter region of *SLC16A6* to promote expression. **(a)** Genome browser tracks of indicated genes of MITF ChIP-Seq from GSE172383: SRX10640872 and SRX10640873. **(b)** RNA sequencing data of *SLC16A6* transcripts with a CRISPR generated *MITF* mutant (\*\**P* < 0.01)^27^. **(c)** As in (b), but with siRNA knockdown of *MITF* (\**P* < 0.05). **(d)** As in (b), but with overexpressed wildtype MITF (\*\*\**P* < 0.001).

**Figure S3.**
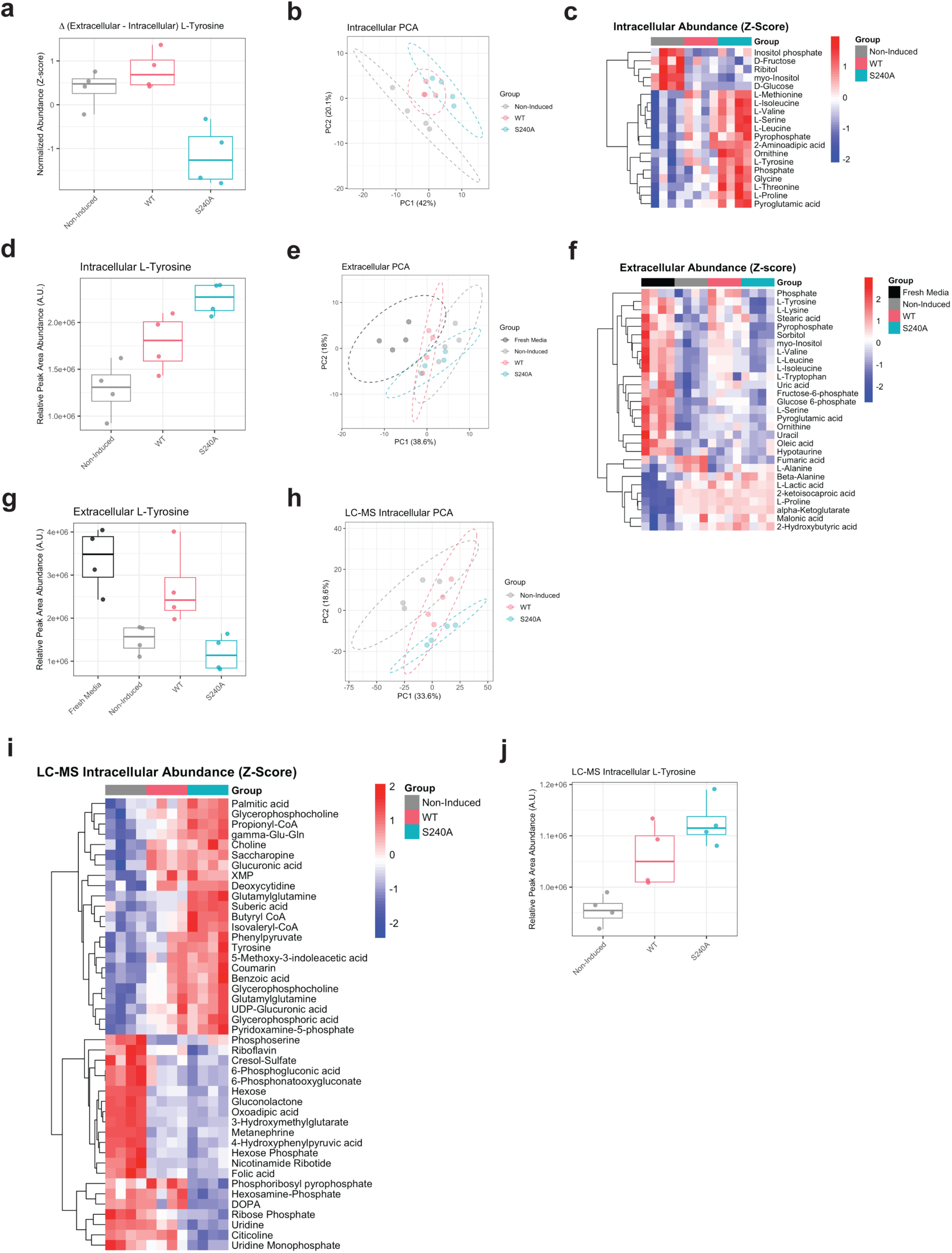
A plasma membrane-rerouted SLC16A6 alters tyrosine-derived metabolites abundances. **(a)** Boxplot of z-scored delta abundance for L-tyrosine measured by GC-MS. **(b-d)** Intracellular GC-MS dataset; **(e-g)** extracellular GC-MS dataset; **(h-j)** intracellular LC-MS dataset. PCA plots (b, e, h) show sample clustering. Heatmaps **(c, f, i)** show metabolites with significantly altered abundance by group. Metabolite abundances were log2-transformed and z-score normalized prior to analysis. Boxplots **(d, g, j)** display relative L-tyrosine abundance measured as relative peak area (a.u.).

**Figure S4.**
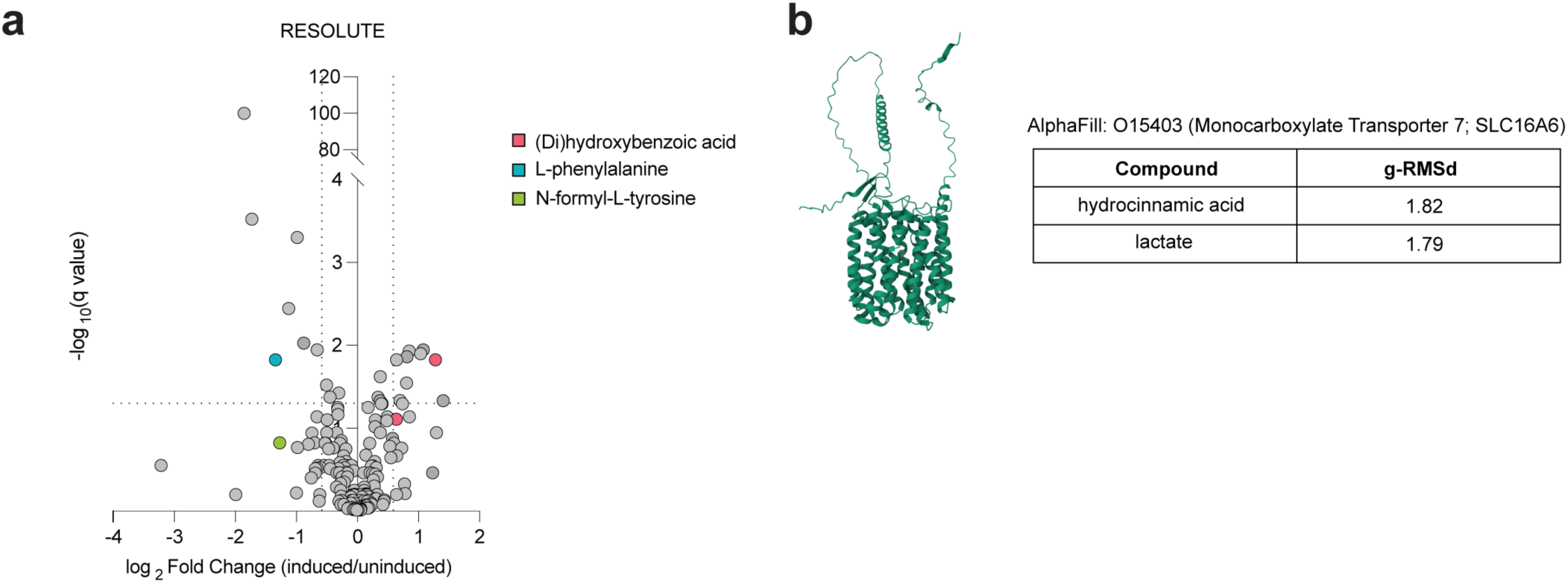
SLC16A6 is predicted to transport aromatic metabolites. **(a)** RESOLUTE metabolomics data of HEK-293T cells induced WT SLC16A6 expression versus non-induced. Data extracted from re-solute.eu/knowledgebase/gene/SLC16A6. **(b)** AlphaFill substrate predictions for SLC16A6. *Left,* structure of SLC16A6 with hydrocinnamic acid. *Right*, table indicating two AlpahFill predicted substrates of SLC16A6.

**Figure S5.**
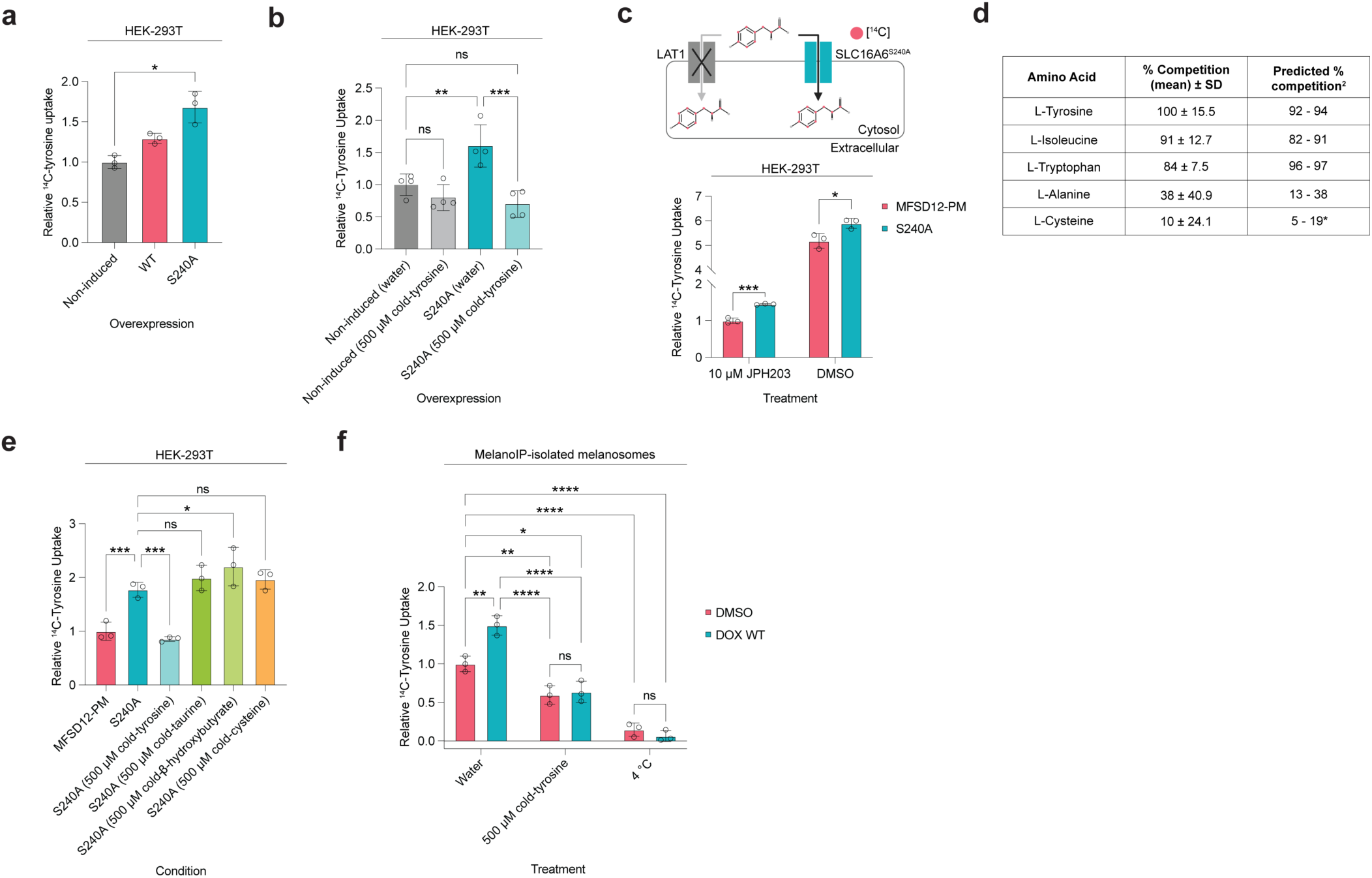
SLC16A6 is sufficient to transport tyrosine. **(a)** Non-induced HEK-293T cells or HEK-293T cells expressing either WT SLC16A6 or SLC16A6^S240A^ were incubated in a buffer containing a LAT1 inhibitor. 5 µM cold-tyrosine and 1 µCi mL^−1 14^C-labeled tyrosine were added to initiate transport. Assay was performed for 10 minutes prior to washes and scintillation. Two-way ANOVA was used for statistical analysis. (\**P* < 0.05; n=3 biological replicates). **(b)** HEK-293T cells non-induced or induced SLC16A6^S240A^-overexpression were incubated in a buffer containing LAT1 inhibitor, JPH203 at 10 µM. Cold-tyrosine (5 µM for transport assay; 500 µM for competition) and ^14^C-labeled tyrosine were added to initiate transport. Assay was performed for 10 minutes prior to washes and scintillation. Two-way ANOVA was used for statistical analysis (\*\**P* < 0.01; n=4 biological replicates). **(c)** HEK-293T cells expressing either MFSD12-PM or SLC16A6^S240A^ were incubated in a buffer containing or lacking 10 µM JPH203. Cold-tyrosine (5 µM for transport assay; 500 µM for competition) and ^14^C-labeled tyrosine were added to initiate transport. Assay was performed for 10 minutes prior to washes and scintillation. Two-way ANOVA was used for statistical analysis (\**P* < 0.05, \*\*\**P* < 0.001, n=3 biological replicates). **(d)** Table comparison between mean percent competition in Fig. 3d versus reported percent competition^2^; asterisk denotes L-cystine competition metrics. **(e)** As in (Fig. 3d), but with cold-taurine, -β-hydroxybutyrate, -cysteine. Two-way ANOVA was used for statistical analysis (\**P* < 0.05, \*\*\**P* < 0.001; n=3 biological replicates). **(f)** As in (Fig. 3e), but with the addition of 500 µM cold-tyrosine or performed at 4°C for 10 min; samples normalized to 1% triton X-100 samples (\**P* < 0.05, \*\**P* < 0.01, \*\*\*\**P* < 0.0001, n=3 biological replicates).

**Figure S6.**
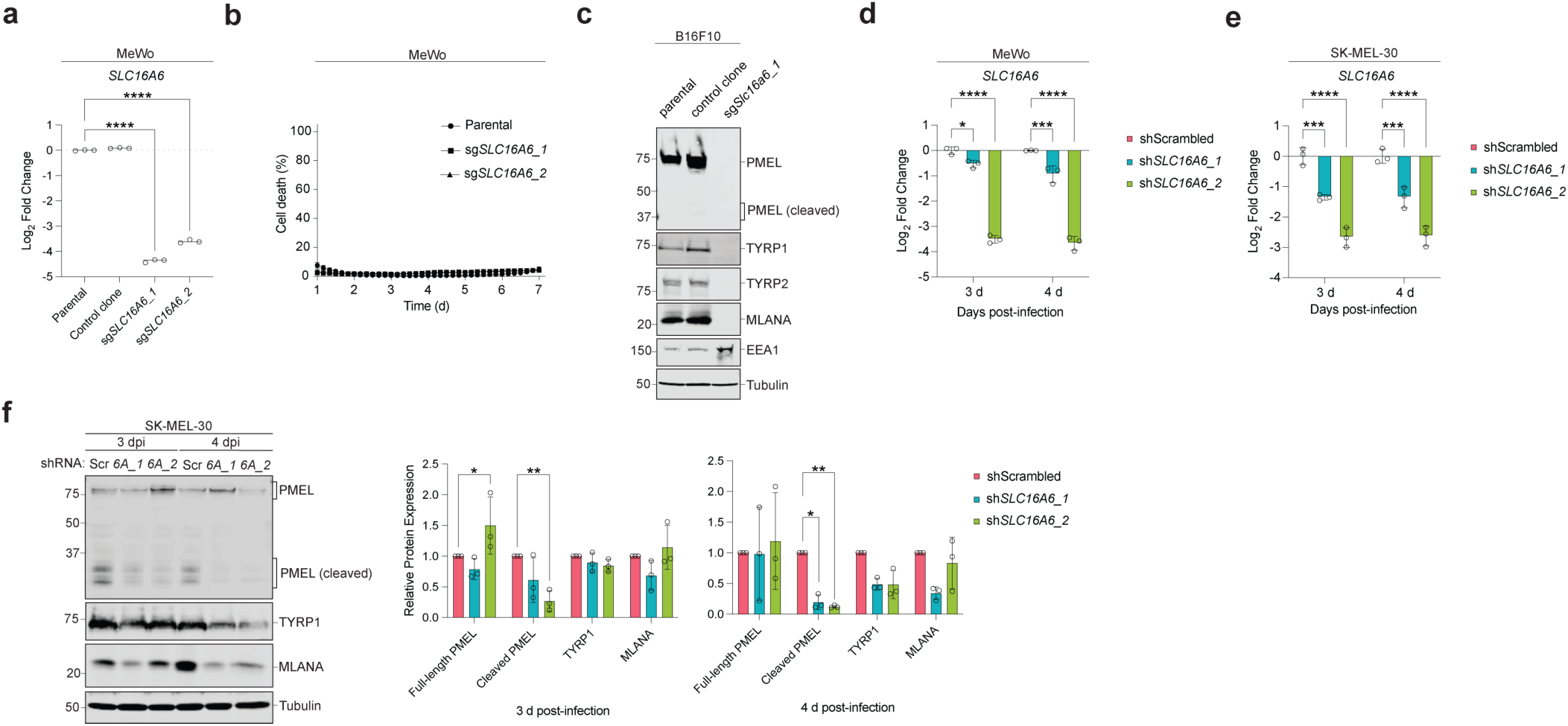
*SLC16A6* depletion downregulates melanosomal components. **(a)** *SLC16A6* transcript abundance changes in parental, control clone, and two *SLC16A6* knockout clones. **(b)** Cell death assay of parental and two *SLC16A6* knockout clones. **(c)** Representative immunoblots of indicated melanogenesis components in B16F10 parental, control clone, and one *SLC16A6* knockout clone (n=3 biological replicates). **(d)** qPCR of *SLC16A6* transcripts in MeWo melanoma cells three and four days post-infection with indicated shRNA lentiviruses. **(e)** qPCR of *SLC16A6* transcripts in SK-MEL-30 melanoma cells three and four days post-infection with indicated shRNA lentiviruses. **(f)** *Left,* representative immunoblots of melanogenesis components in SK-MEL-30 cells three and four days post-infection with indicated shRNA lentiviruses (*Scr*, scrambled shRNA control; *A6_1*, shRNA targeting SLC16A6 #1; *A6_2*, shRNA targeting SLC16A6 #2; n=3, biological replicates). *Right*, quantification of relative protein expression of shRNA treatment in at least n=3 biological replicates. Each protein is normalized to control alpha-tubulin (\**P* < 0.05; \*\**P* < 0.01).

